# Pneumo-Typer: integrated genomic surveillance tool for capsule genotype, serotype and sequence type in *Streptococcus pneumoniae* informs vaccine strategies

**DOI:** 10.1101/2025.02.13.638184

**Authors:** Xiangyang Li, Huejie Zhang, Yaoyao Zhu, Zilin Yang, Xiangyu Wang, Guohui Zhang, Xuan Zhao, Yinyan Huang, Bingqing Li, Zhongrui Ma

## Abstract

**Background:** *Streptococcus pneumoniae* persists as a leading global pathogen, with current capsule-based vaccines like PCV13 offering limited coverage against its extensive serotype diversity (>100 serotypes). This restricted coverage drives the emergence of non-vaccine serotypes. Moreover, vaccine efficacy is further compromised by immune evasion caused by capsule genotype (CapT) variations within vaccine-covered serotypes, while existing genome-based tools lack robust analytical capabilities for CapT.

**Results:** We developed Pneumo-Typer, a CapT visualization tool enabling rapid identification of CapT variations in *S. pneumoniae*. Application to 17,407 publicly available genomes from the NCBI database revealed widespread CapT variations across vaccine-covered serotypes, providing critical insights for optimizing vaccine formulations. In addition, Pneumo-Typer integrates high-accuracy serotype prediction (98.44% accuracy across 93 serotypes) and multilocus sequence typing (ST), facilitating comprehensive genomic surveillance. Besides, CapT and ST information can assist in serotype determination. Pneumo-Typer enables integrated analysis of serotype-CapT-ST relationships, demonstrating that while most serotypes associate with multiple STs and CapTs, ST and CapT lineages evolve independently. Strikingly, multi-serotype serogroups exhibited pronounced CapT diversity, suggesting evolutionary processes driving serogroup expansion through CapT plasticity.

**Conclusions:** In summary, Pneumo-Typer is an integrated platform for genomic surveillance of pneumococcal populations. By resolving previously underexplored serotype-CapT-ST linkages, the tool provides a framework for next-generation vaccine design, emphasizing the need for vaccine formulations that address both serotype replacement and CapT-driven immune evasion.

## Background

*Streptococcus pneumoniae* remains a leading global pathogen responsible for over 1.2 million annual deaths due to pneumonia, meningitis, and invasive pneumococcal disease (IPD), particularly among young children, the elderly, and immunocompromised individuals [1, 2]. The polysaccharide capsule, which defines over 100 distinct serotypes, is both a critical virulence factor and the primary target of pneumococcal vaccines [3, 4]. Current vaccines, such as the 13-valent pneumococcal conjugate vaccine (PCV13), have significantly reduced disease burden by targeting predominant serotypes [5].

However, pneumococcal vaccine efficacy is increasingly challenged by two phenomena: serotype replacement driven by the expansion of non-vaccine serotypes [6–8], and capsular genotype (CapT)-driven immune evasion within vaccine-targeted serotypes. While serotype replacement has been widely recognized and addressed through continuous updates to vaccine serotype composition [9], CapT variations remain an emerging concern. These genetic alterations in capsule biosynthesis can lead to antigenic shifts, allowing strains to evade vaccine-induced protection [10–13]. For instance, several strains classified as serotype 14 by molecular-based serotyping tools (SeroBA and SeroCall) exhibit significant alterations in their CapTs, allowing them to bypass vaccine protection [14]. Notably, these molecular methods may fail to fully capture CapT variations, further complicating surveillance efforts.

Traditional serotyping methods, such as the Quellung reaction, are labor-intensive and require specialized expertise [15–17]. Molecular-based serotyping tools, including PneumoCaT [18] and SeroCall [19], utilize sequence alignment approaches but depend on high-quality whole genome sequencing (WGS) data. Other tools, such as SeroBA [20], PneumoKITy [21], and PfaSTer [22], employ *k*-mer-based approaches for faster serotyping but may struggle with mosaic sequences, which are prevalent in pneumococcal populations [23]. Detecting and analyzing CapT variations remains particularly challenging due to the complexity of current bioinformatics workflows, which often require multiple platforms and substantial computational expertise [24, 25]. These challenges underscore the urgent need for high-resolution molecular surveillance tools capable of tracking both serotype dynamics and CapT variations.

While WGS has improved resolution for lineage tracking through sequence types (STs) [26] or global pneumococcal sequence clusters (GPSCs) [27], existing bioinformatics tools lack integrated analysis of capsular locus heterogeneity (CapT variations), serotype prediction, and ST assignment in a unified framework [28]. This gap hinders comprehensive surveillance of vaccine-driven selection pressures, as exemplified by the emergence of serotype 19A-ST320 [29], serotype 8-ST53 [30], and serotype 3-ST271 [31].

To address these challenges, we developed Pneumo-Typer, a high-throughput analytical pipeline that enables simultaneous identification of CapTs, serotypes, and STs (with an optional cgST) from assembled WGS data. The tool demonstrated high efficiency by processing 21,260 genomes within approximately one hour while achieving 98.44% serotyping accuracy across 93 serotypes. Application to 17,407 publicly available pneumococcal genomes from the NCBI database revealed extensive CapT variations, providing critical insights for optimizing vaccine formulations. Furthermore, systematic analysis elucidated complex CapT-serotype-ST relationships, revealing previously unrecognized patterns of pneumococcal evolution. This integrated tool bridges the gap between genomic surveillance and vaccine development, offering a scalable solution for monitoring pneumococcal evolution in the post-pneumococcal vaccine era.

## Implementation

### Pneumo-Typer workflow

Pneumo-Typer, developed in Perl, offers a distinct advantage over several available Python-based serotyping tools due to its smooth installation on UNIX platforms such as Linux and macOS. The tool requires specific Perl modules, including GD and GD::SVG, as well as BLAST+, BLAT, and Prodigal. For user convenience, Pneumo-Typer is also available as a Bioconda package with preinstalled dependencies. As shown in Fig. 1, the workflow of Pneumo-Typer is divided into three main steps:

**Fig. 1.**
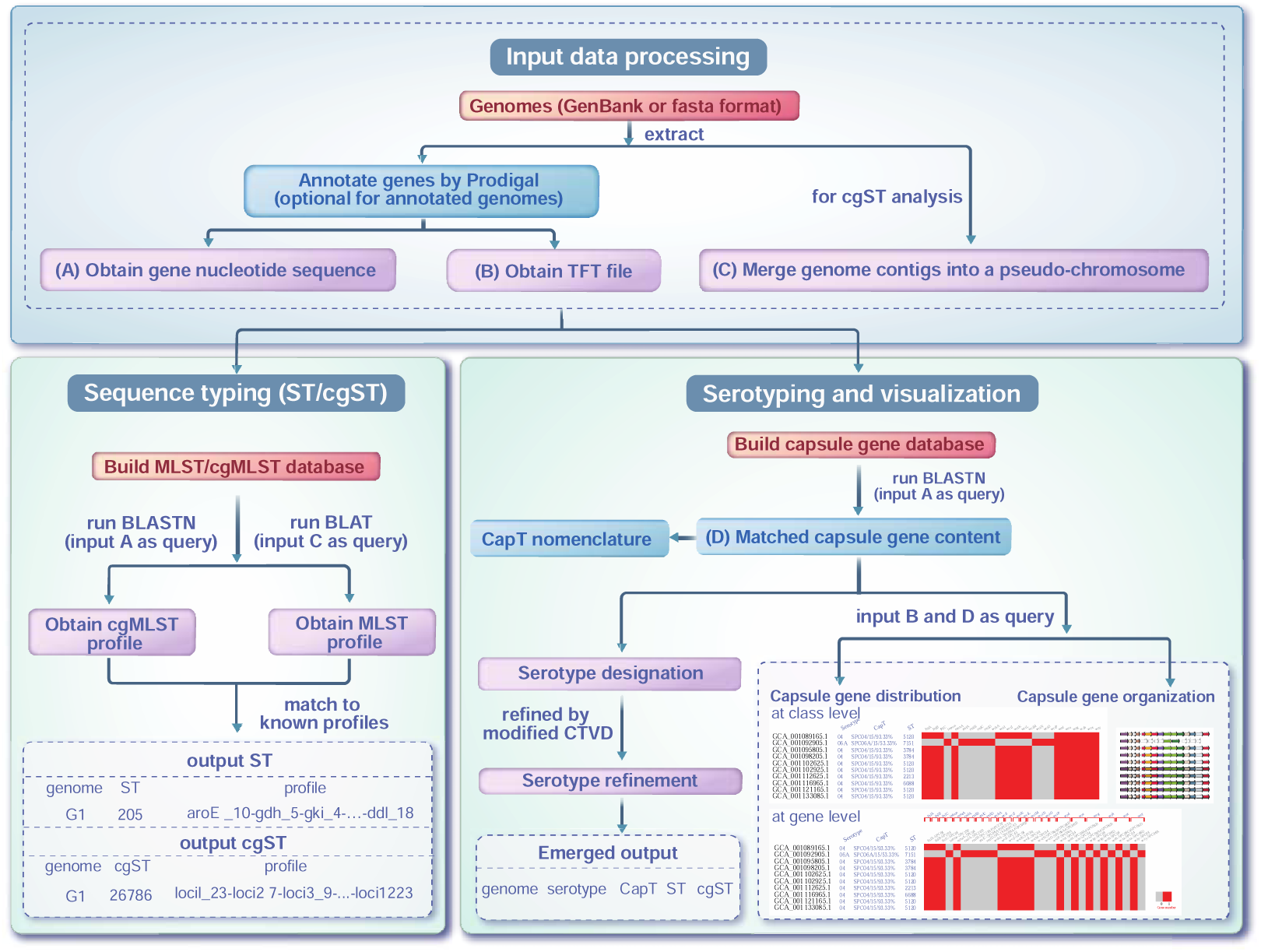
The workflow of Pneumo-Typer.

#### Step 1: Input data processing

Genomes provided in GenBank or FASTA formats undergo systematic preparation. For each genome, gene nucleotide sequences (A) and TFT-formatted feature tables (B) - seven-column tab-delimited files detailing genomic features - are initially extracted [32]. Genomes lacking existing annotations are processed using Prodigal for automated gene prediction [33]. To ensure methodological consistency across datasets, Prodigal may optionally be applied to all input genomes regardless of their initial annotation status. For subsequent cgMLST (core genome multilocus sequence typing) analysis, a critical preprocessing step involves the seamless concatenation of all genomic contigs into a unified pseudo-chromosome (C), effectively simulating complete genome architecture for downstream comparative analyses.

#### Step 2: Sequence typing

##### MLST/cgMLST database retrieval

For MLST, seven allele genes (*aroE*, *gdh*, *gki*, *recP*, *spi*, *xpt*, *ddl*) along with their related allelic profiles specific to pneumococcus were sourced from the PubMLST website (https://pubmlst.org/) [34]. In parallel, for cgMLST, sequences from a collection of 1,222 allele genes and their matching allelic profiles were acquired for the determination of the cgST.

*ST and cgST determination*. Inputs A (gene sequences) or C (pseudo-chromosome) served as queries for allele identification. The ST determination was executed through a systematic approach: (i) The genes were compared with the MLST database using BLASTN. (ii) Stringent criteria (E-value = 1e^−5^, identity = 100%, coverage = 100%) were applied on the BLASTN results to identify multilocus sequence gene distribution. (iii) Identified genes were unified using dashes, resulting in a profile like aroE_13-gdh_13-gki_41-recP_5-spi_13-xpt_1-ddl_31. Missing allele genes in the genome were represented as “X”. (iv) STs were determined by matching profiles against known ST allelic profiles. Genomes with an “X” got an unknown ST label, and those with novel profiles were tagged as novel STs. The cgST determination followed a similar blueprint but with nuances: (i) BLAT [35] was harnessed for precise mapping of cgMLST loci against genomes. (ii) The “-fastMap” parameter sped up the process, fusing 1,222 cgMLST sequences into one, but for sequences like SPNE00213.fas that exceeded 5,000 bp, “-fastMap” was not used. (iii) Missing allele genes in the genome get a unique marker, like SPNE00001_N.

#### Step 3: Serotyping and visualization

##### Capsule gene database construction

The capsule gene database was established by collating GenBank files associated with capsule gene loci from reference strains, which covered a spectrum of 93 serotypes. The foundation of the database, comprising 90 serotypes, was extracted from the EMBL Nucleotide Sequence Database under accession numbers CR931632–CR931722 [36]. The dataset was enriched with two more serotypes 6C and 35D, sourced from NCBI under accession numbers JF911515 [37] and KY084476 [38], respectively. Serotype 6D’s sequence was procured from the PneumoCaT database [18] and annotated leveraging Prokka v1.12 [39]. 6E and 23B1 were not included as independent serotypes in the capsule gene database since they emerged as novel CapT subtypes. For instance, the isolate carrying the “6E” capsule gene locus yielded the serotype 6B capsule [40, 41]. The nucleotide sequences of the 93 serotypes were extracted from the GenBank files by a custom Perl script. Subsequently, each capsule gene was tagged with its respective gene name and serotype, exemplified as *wzd*-SPC30. The genes underwent a two-step refinement: (i) exclusion of genes with incomplete sequences or encoding transposases. (ii) dereplication of genes sharing 100% nucleotide identity using a proprietary Perl script. Upon completion, the capsule gene database encompassed 1,290 genes spread across the 93 serotypes. The database included two principal files: a sequence file and a list file. The sequence file, formatted in fasta, delineated sequence IDs preceding their respective sequence data. Every sequence ID included a unique ID complemented by gene annotation, separated by a whitespace (e.g., SPC16F_0021 dTDP-D-glucose_4,6-dehydratase_RmlB). The list file, structured in a tab-delimited format, comprised two columns. The first column denoted the gene class and its abbreviation, distinctly separated by dual underscores. The second column hosted the unique ID for each gene, with a comma serving as a delimiter for multiple genes when required (e.g., wzy_wzy-SPC24B [’SPC24B_0019’, ‘SPC24B_0015’]).

##### Serotype determination

Serotype determination was performed by querying Input A (gene sequences) against our established capsule gene database using BLASTN, with matches filtered by stringent thresholds (identity ≥70%, coverage ≥95%, E-value ≤1e−5). Genes aligning to multiple serotypes were counted across all relevant categories, and the predominant serotype was assigned based on the highest cumulative gene matches (called Serotype Designation). Ambiguities were resolved using serotype-specific markers from a modified Capsular Typing Variant Database (CTVD) (called Serotype Refinement) (Table S1).

##### CapT nomenclature and visualization

We established a CapT nomenclature system. For example, the CapT of strain GCA_001457635.1 was designated as SPC01/15/93.33%, where “SPC01” indicates serotype 1, “15” represents the number of matched capsule genes, and “93.33%” signifies that 14 out of 15 matched capsule genes belong to serotype 1. For strains that could not be distinguished using these three criteria, numerical suffixes were added for further differentiation (e.g., SPC04/16/81.25%-1 and SPC04/16/81.25%-2), ensuring that each CapT corresponds to a unique capsule gene content. For CapT visualization, capsule gene distribution and operon organization were mapped using Inputs B (TFT files) and D (capsule gene content). Genes affected by frameshift mutations, nonsense mutations, or terminator codon mutations were not displayed in any CapT visualization (Table S2). A custom-modified version of Gcluster [32] generated two-tiered heatmaps (gene- and class-level resolution) from tab-delimited positional data, while TFT-derived annotations enabled operon structure visualization. CapT, Serotype and ST/cgST metadata were algorithmically superimposed onto these graphical outputs.

### Data acquisition, processing and analysis

#### Data acquisition and processing

To test the performance of Pneumo-Typer, all pneumococcal genomes from the NCBI database were downloaded as of Oct 5, 2022. These genomes, procured in GenBank format, were extracted using a custom Perl script aligned with a CSV file input. This CSV file was readily available from the NCBI genome database website (https://www.ncbi.nlm.nih.gov/genome/browse/) by searching with the term “*Streptococcus pneumoniae*”. Furthermore, the serotype information was retrieved from the GenBank files through our proprietary Perl program. To ensure the quality and accuracy of the genomes, two diagnostic tools: CheckM [42] and QUAST [43] were employed. CheckM evaluated each genome’s completeness and potential contamination based on lineage-specific sets of single-copy genes. Genomes boasting near-perfection (≥90% completeness) and negligible contamination (≤5% contamination) [44] were selected. QUAST offered metrics on assembly quality, and the genomes were sieved by stringent criteria, including contig size ≤ 500 bp and N50 ≥ 40 kb.

#### Data analysis

As shown in Fig. 2, after filtering process, 21,260 genomes were obtained, and used to assess the performance of Pneumo-Typer in ST and cgST determination. Among these, 17,667 genomes with recorded serotypes (covering 82 serotypes) were selected and combined with 16 genomes (covering 11 serotypes) downloaded from an earlier investigation [18] and assembled via MetaHIT [45], yielding a total of 17,683 genomes (covering 93 serotypes) for evaluating Pneumo-Typer’s performance in serotype determination. Among these, 17,407 genomes that were accurately serotyped by Pneumo-Typer were used to analyze the relationships between serotypes, STs, and CapTs. Among these, 14,716 genomes within vaccine-covered serotypes were selected to evaluate Pneumo-Typer’s performance in CapT analysis.

**Fig. 2.**
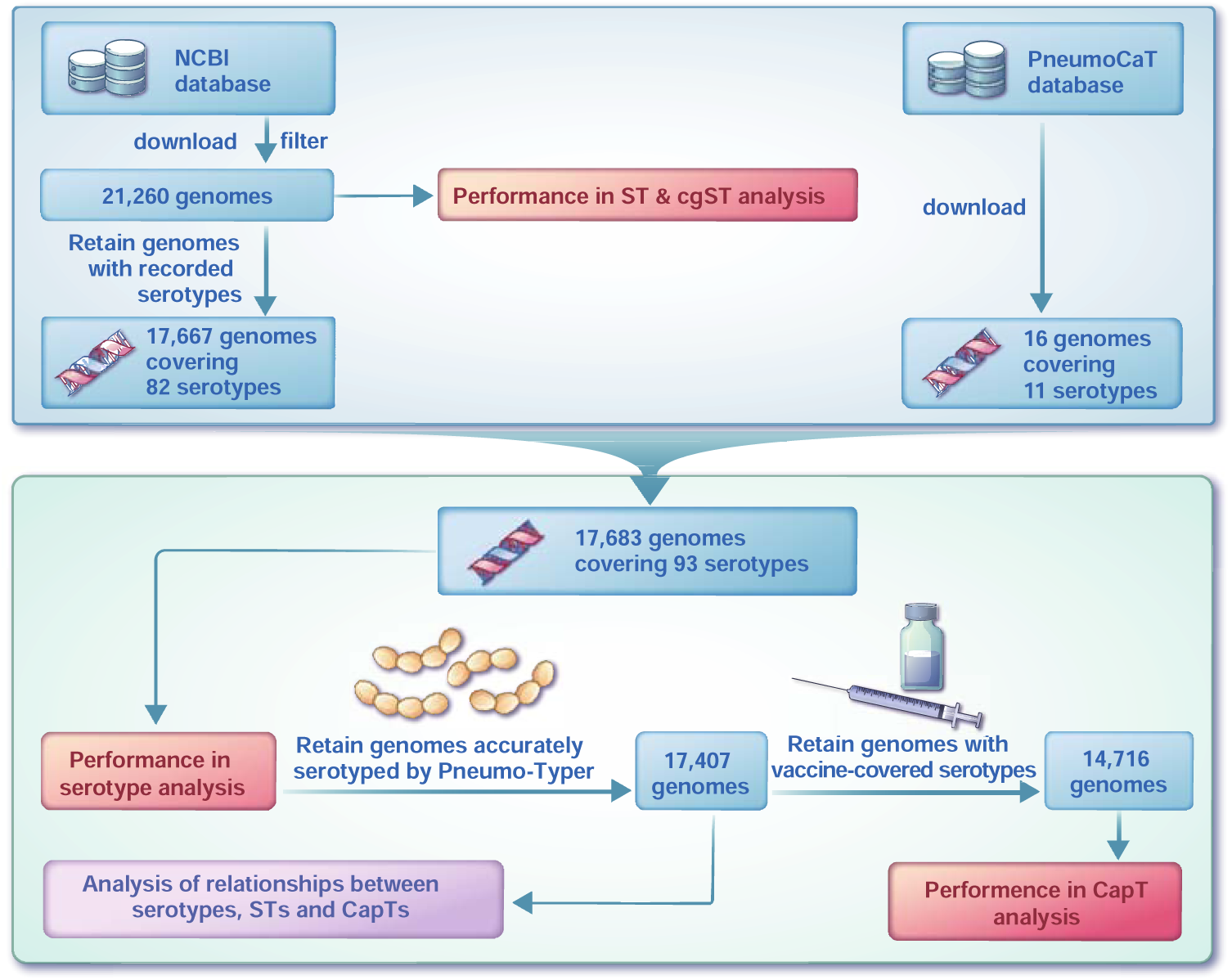
Data acquisition, processing, and analysis workflow for this study.

The analysis was performed on an Ubuntu 20.04 server equipped with four Intel Xeon Platinum 8260L 24-core processors and 512 GB RAM, harnessing 192 CPU threads. Parameter -p was set to “T”, prompting Prodigal to annotate all genomes under examination during Pneumo-Typer operation.

#### Clonal Complexes analysis

The concept of MLST Clonal Complexes (CC) refers to STs that have differences in a single locus variant (SLV) based on the eBURST algorithm[46]. The analysis was performed using the PHYLOViZ 2.0 software[47].

#### ST or CapT diversity calculation

In this study, Shannon’s entropy was employed to assess the diversity of STs or CapTs within individual serotypes. Shannon’s entropy is equal to:

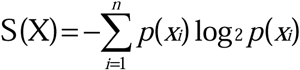

Where S(X) represents the Shannon’s entropy of the variable X (either ST or CapT), *p*(*x_i_*) denotes the probability of X taking the *i*-th value *x_i_*, and *n* is the total number of possible values for X.

#### Statistical analysis

Spearman’s correlation was used to analyze the relationship between two sets of variables. A correlation coefficient |R| ≥ 0.8 indicates a strong correlation, while 0.3 ≤ |R| < 0.5 suggests a weak correlation, and |R| < 0.3 denotes little to no correlation. To assess statistical differences between the two groups, an independent samples *t*-test was performed, with significance indicated by * *P* < 0.05.

## Results

### Key features of Pneumo-Typer

Pneumo-Typer has four key features:

#### Simple Input

The only mandatory input for Pneumo-Typer is a directory containing genomes in either GenBank or fasta format, or a combination of both, ensuring straightforward acquisition of input data.

#### Batch Processing

Pneumo-Typer efficiently handles the serotype, ST, cgST (optional), and CapT of a large array of pneumococcal genomes simultaneously using multiple threads. As a benchmark, it processed 21,260 genomes in approximately 53 minutes under default settings, with all genomes annotated using Prodigal and subjected to serotype and ST analysis, utilizing 190 threads on an Ubuntu 20.04 server powered by four Intel Xeon Platinum 8260L processors.

#### Innovative Serotyping

Pneumo-Typer employs a novel two-step sequence alignment approach for serotyping. The first step, termed Serotype Designation, involves BLASTing genomes against a custom capsule gene database (covering 93 dominant serotypes). Serotype is assigned based on the serotype(s) with the highest proportion of matched genes. The second step, known as Serotype Refinement, further refines the serotype using serotype-specific markers from a modified CTVD (Table S1). If the serotypes predicted in Steps 1 and 2 do not belong to the same serogroup, the serotype from Step 1 is retained to avoid errors due to mosaic sequences.

#### Advanced CapT Visualization

Pneumo-Typer generates SVG-formatted heatmaps that display the capsule gene distribution at the gene (heatmap_gene.svg) or class (heatmap_class.svg) level. Additionally, it creates an SVG-formatted map that graphically represents the capsule operon organization (CPS_operon.svg), with distinct homologous genes depicted in varied colors. To enable superior comparison, there is also an option to map the capsule contexts onto a user-provided phylogenetic tree.

### Performance of Pneumo-Typer in serotyping and sequence typing

At the serogroup level, Pneumo-Typer reached 99.84% accuracy (Table S3). At the serotype level, Pneumo-Typer achieved an accuracy of 84.98% (15,027 out of 17,683) after step Serotype Designation, which increased to 98.44% (17,407 out of 17,683) following step Serotype Refinement (Table 1). Herein, Pneumo-Typer achieved 100% prediction accuracy for 55 serotypes, 99%-100% accuracy for 10 serotypes, and 98%-99% accuracy for 7 serotypes. However, Pneumo-Typer had low prediction accuracy for some serotypes, such as 12A (0%), 25A (0%), 32A (0%), 44 (0%), 9A (20%), and 35C (31.48%). For comparison, we tested PneumoKITy and PfaSTer on the same dataset. PneumoKITy achieved an accuracy of 69.01% (12,203 out of 17,683), and PfaSTer achieved 90.29% (15,966 out of 17,683) (Table S4).

**Table 1.** The performance of Pneumo-Typer in serotyping.

| No. | Recorded serotype (S) | Serotype Designation (D) | Serotype Refinement (R) | Number | Percentage (%) | D = S | R = S |
| --- | --- | --- | --- | --- | --- | --- | --- |
| 1 | A | A | A | 14646 | 82.83 | yes | yes |
| 2 | A | B | A | 2410 | 13.63 | no | yes |
| 3 | A/B | A | A | 272 | 1.54 | yes | yes |
| 4 | A/B | A/B | A | 79 | 0.45 | yes | yes |
| 7 | A | B | B | 211 | 1.19 | no | no |
| 5 | A | A | B | 30 | 0.17 | yes | no |
| 6 | A | A/B | A/B | 24 | 0.14 | no | no |
| 8 | A | A/B | B | 8 | 0.05 | no | no |
| 9 | A | B/C | B/C | 3 | 0.02 | no | no |
| **Total** |  |  |  | **17683** | **100.00** | **Accuracy:**<br>**84.98%** | **Accuracy:**<br>**98.44%** |

**Notes:** A, B, A/B, and B/C indicate possible serotype scenarios. Recorded serotypes refer to the serotypes documented in the NCBI Database. The definitions of Serotype Designation and Serotype Refinement are detailed in the "Overview of Pneumo-Typer" section.

The CapT visualization features of Pneumo-Typer allowed us to specifically examine the CapTs of the mispredicted strains by Pneumo-Typer. Herein, a strain recorded as serotype 1 but predicted as 16F had a capsule operon organization distinctly resembling 16F (Fig. 3A). Similarly, two strains recorded as serotype 6D but predicted as 6B showed a capsule operon organization more consistent with 6B, as 6B had a gap between the *wciN* and *wciO* genes, which 6D lacked (Fig. 3B). Additionally, four strains recorded as serotype 7B but predicted as 7C had a capsule operon organization that matched both 7B and 7C (Fig. 3C), yet their capsule gene distribution at the gene level aligned more closely with 7C (Fig. 3D). Excluding these misannotated serotypes would further improve the serotype prediction accuracy of Pneumo-Typer. After exclusion, the accuracy for serotype 1 would increase from 99.90% to 100%, for serotype 6D from 84.62% to 100%, and for serotype 7B from 86.67% to 100%. Therefore, Pneumo-Typer can not only predict serotypes but also verify the correctness of annotated serotypes through its CapT visualization features.

**Fig. 3.**
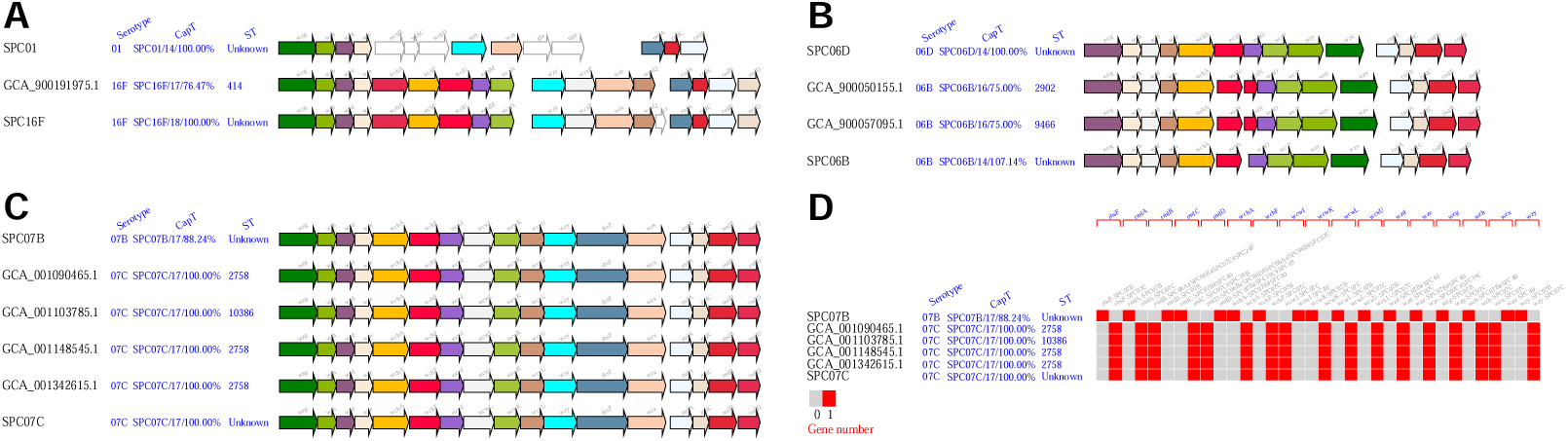
The role of CapT visualization in serotyping. **A** CPS operon map of serotype 1 reference strain, GCA_900191975.1, and serotype 16F reference strain. **B** CPS operon map of serotype 6D reference strain, GCA_900050155.1, GCA_900057095.1, and serotype 6B reference strain. **C** CPS operon and **D** heatmap of serotype 7B reference strain, GCA_001090465.1, GCA_001103785.1, GCA_001148545.1, GCA_001342615.1, and serotype 7C reference strain.

Pneumo-Typer demonstrated exceptional performance in the rapid assignment of STs, processing 21,260 genomes in under 15 minutes. The assigned STs revealed the presence of 3,250 known STs and the emergence of 18 novel STs (Table S5). However, 311 genomes remained unassigned to any ST, likely due to incomplete ST locus identification in draft-status genomes. Among the identified STs, several exhibited high isolate counts, notably ST217 (459 isolates), ST156 (390 isolates), ST180 (387 isolates), ST199 (349 isolates), and ST191 (319 isolates). A substantial number of STs, 1,801, were assigned to only one isolate each (Table S6).

Besides, using Pneumo-Typer, 10,859 out of 21,260 genomes were categorized into 10,103 distinct cgSTs (Table S5). Additionally, the analysis revealed 10 novel cgST profiles. The most frequently occurring cgST was cgST12235, accounting for 141 isolates, followed by cgST1494, which accounted only for 7 isolates. A substantial number of cgSTs, 9,600, were assigned to only one isolate each. However, a significant number of genomes, 10,391, could not be linked to established cgSTs, indicating that known cgSTs could only be attributed to almost half of the genomes. This suggests a limitation in using cgST for molecular typing, given the extensive number of gene loci required, which may not be fully assembled in draft-status genomes.

### Performance of Pneumo-Typer in identifying CapT variations

Serotypes represent phenotypic characteristics, whereas CapT denotes the genotype, with the latter determining the former. However, due to the limitations of molecular-based serotyping tools, the serotype results obtained may not always accurately reflect CapT variations, which can significantly impact the identification of potential capsule antigenic shifts. To address this issue, we employed CapT visualization features generated by Pneumo-Typer and evaluated its performance using three serotype 14 variants reported in a previous study firstly [14]. WGS data from these three serotype 14 variants (PMP1437, PMP1438, and PMP1514), along with the serotype 14 reference strain (SPC14), were analyzed using Pneumo-Typer, resulting in distinct types of CapT profiles (Fig. 4). Although Pneumo-Typer misclassified all three isolates as serotype 14 or 13|20, its CapT visualization features revealed critical genetic differences. Specifically, PMP1437, PMP1438, and PMP1514 all exhibited multiple gene deletions (Fig. 4B). Additionally, PMP1514 contained an inserted sequence that did not match any known genes in our capsule gene database (Fig. 4A). Moreover, PMP1437 appeared to be a hybrid, incorporating genes from both serotype 13|20 and serotype 14 (Fig. 4C). These findings demonstrate that Pneumo-Typer’s CapT visualization features effectively identify CapT variations, providing valuable insights into genetic changes that may not be captured by conventional serotyping methods.

**Fig. 4.**
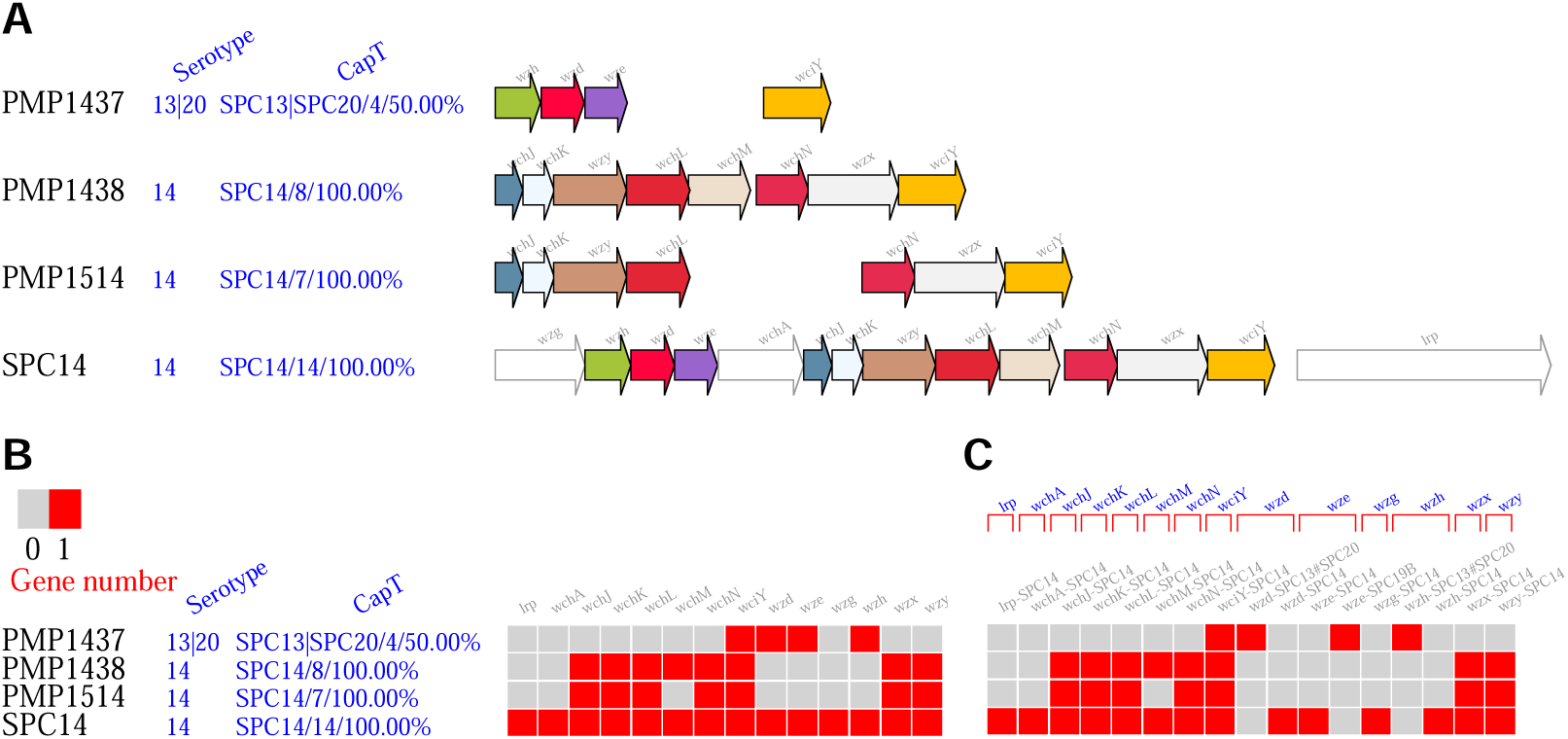
CapT visualization of three serotype 14 variants by Pneumo-Typer. **A** Capsule operon organization. **B** Capsule gene distribution at class level. **C** Capsule gene distribution at the gene level. Serotype 14 variants: PMP1437, PMP1438, and PMP1514; Serotype 14 reference strain: SPC14. Any genomic region that does not match the capsule gene database appears as a gap in the Capsule operon organization diagram.

In the context of vaccine design, monitoring capsule antigenic shifts in vaccine-covered serotypes is crucial, as these shifts directly impact vaccine efficacy. To address this issue, we analyzed the CapT profiles of serotypes included in PPSV23, PCV13, PCV15, PCV20, PCV21, and PCV24 [48] to identify CapT variations, including gene mutations and horizontal gene transfer events (Table 2). Notably, several CapT variations with high prevalence warrant further attention (Fig. 5). For serotype 14, mutations were identified in the *wciY* gene, which encodes a putative glycerol phosphotransferase, in 1,232 out of 1,412 strains (87.1%). However, previous studies have shown that the disruption of *wciY* does not impact capsular production [49]. The *rmlD* gene, which encodes dTDP-4-dehydrorhamnose reductase, was mutated in 648 out of 1,269 serotype 19A strains (51.1%) and 75 out of 248 serotype 23B strains (30.2%). Given that *rmlD* is essential for L-rhamnose biosynthesis in the capsule, its disruption is expected to influence capsular structure. For serotype 11A, mutations were detected in the *wcwC* gene, encoding an O-acetyltransferase, in 63 out of 402 strains (15.7%). In serotype 20, mutations in the *whaF* gene were identified in 91 out of 111 strains (82.0%). Notably, the *whaF* mutant has been designated as serotype 20A, whereas mutations in *wciG* (detected in 4 out of 111 strains) have led to the classification of serotype 20C. These variants are distinct from serotype 20B (which corresponds to serotype 20 in this study), with 20A and 20B included in PCV21 and PCV24, respectively [50, 51]. Additionally, in serotype 33F, an insertion of the *wcyO* gene, encoding a putative acetyltransferase from serotype 33C, was observed in 23 out of 145 strains (15.9%). Similarly, serotype 17F exhibited an insertion of the *glf* gene, encoding UDP-galactopyranose mutase, likely acquired from serotype 33B, in 49 out of 154 strains (31.8%). Given the role of *glf* in the biosynthesis of UDP-galactofuranose (Gal*f*) [52], the impact of this insertion on the capsule structure of serotype 17F remains unclear. These findings highlight the potential risk that vaccines based on standard serotype strains may lose efficacy or fail to protect against emerging CapT variants. Integrating these CapT variants into future vaccine formulations may serve as a viable strategy to maintain or enhance vaccine coverage. Moreover, Pneumo-Typer proved to be an effective tool for serotype subtyping, as demonstrated in this study by its ability to differentiate between the serotype 20 subtypes 20A, 20B, and 20C.

**Fig. 5.**
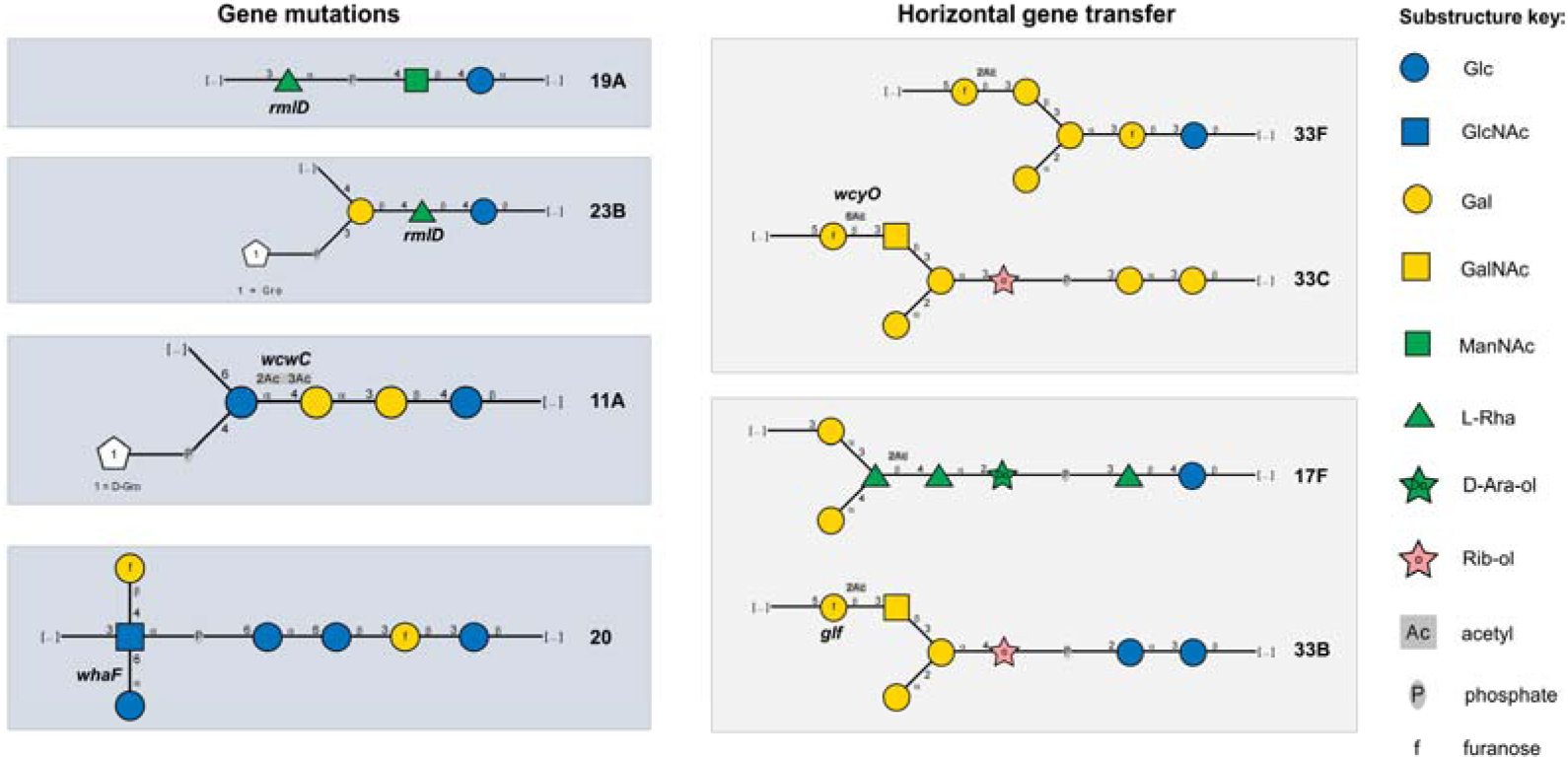
Capsule repeating unit of several serotypes with CapT variations in this study.

**Table 2.**
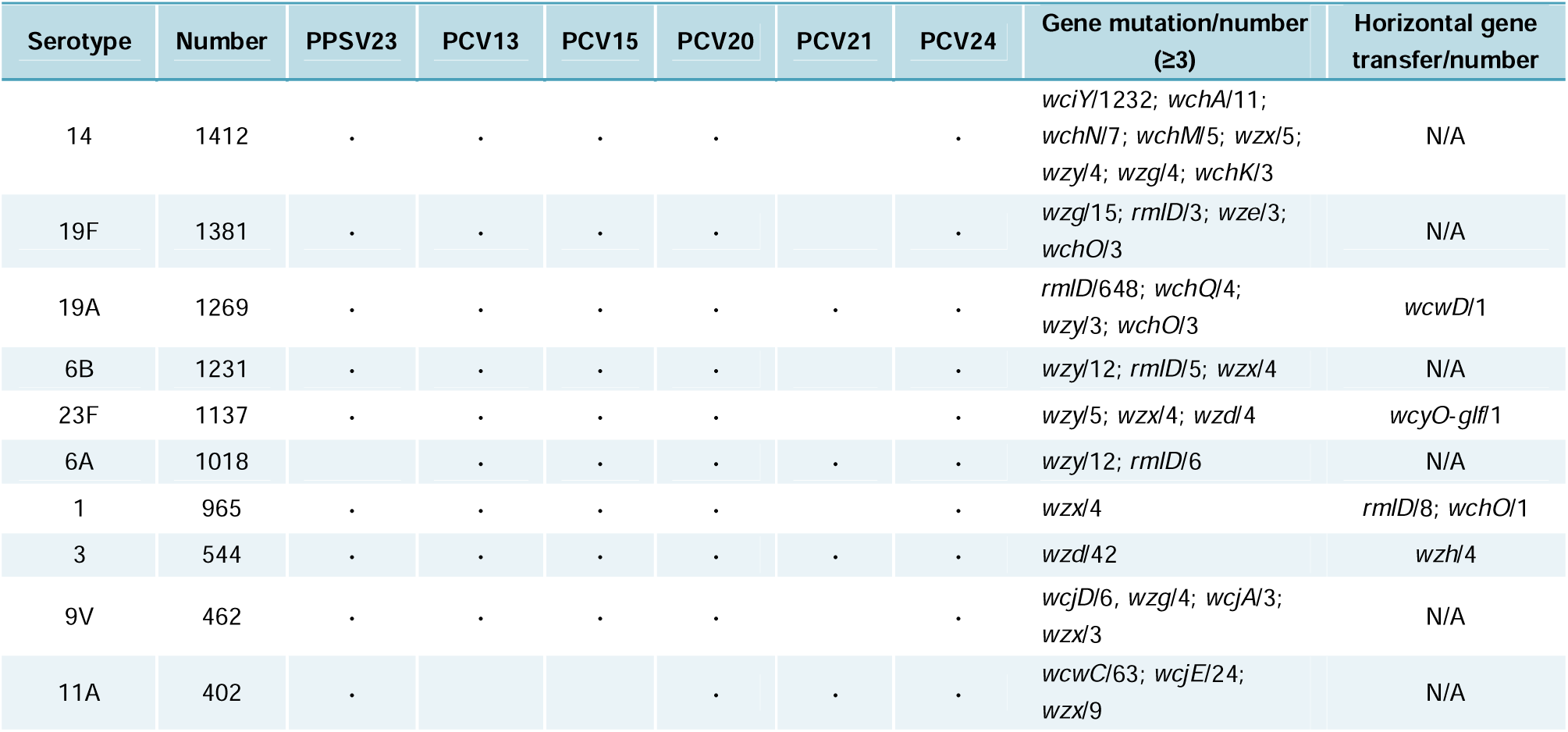
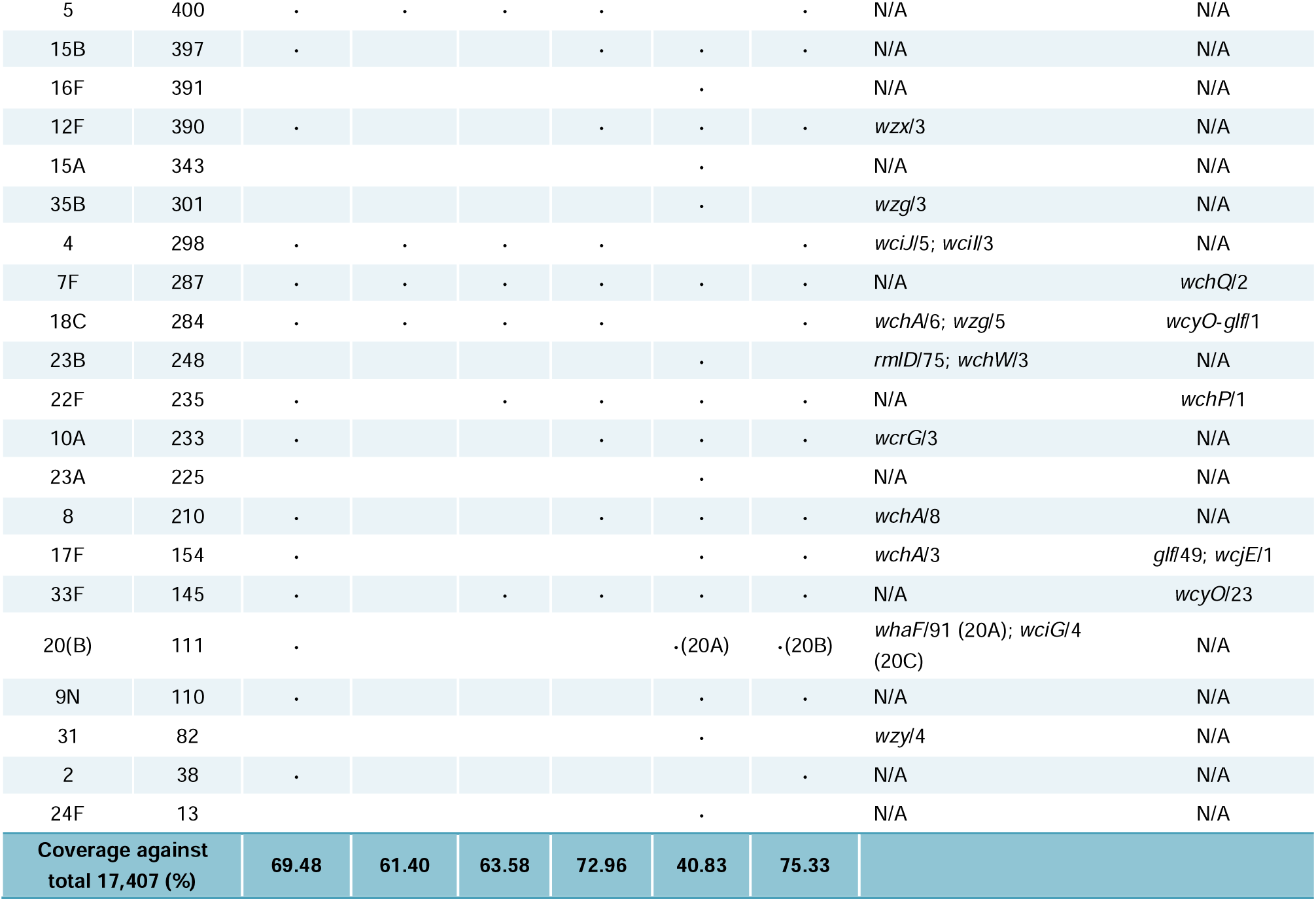
Key CapT variations in vaccine-targeted serotypes.

### Bidirectional serotype-ST associations

The performance of Pneumo-Typer in both serotyping and sequence typing enables a systematic study elucidating the relationships between serotypes and STs. The relationships between serotypes and STs (Table S7) can be analyzed in two ways: the STs associated with each serotype and the serotypes associated with each ST.

First, we analyzed the STs associated with each serotype (Table S8) and found that the vast majority—78 out of 90 serotypes—were associated with multiple STs (Fig. 6A). A subset of serotypes, specifically 19F (≥284 STs), 6B (≥257 STs), 6A (≥238 STs), 19A (≥216 STs), and 23F (≥182 STs), exhibited a particularly high number of coexisting STs. In contrast, a small group of serotypes, including 7A, 10C, 11C, 11D, 11F, 16A, 19C, 24B, 42, 43, 47A, and 47F were observed to have only one unique ST association. However, the ST richness of each serotype heavily depended on its sample size (Fig. 6B). To evaluate the ST diversity of each serotype independently of sample size, we excluded serotypes with fewer than 100 isolates to avoid potential bias and used Shannon’s entropy as a measure of diversity (higher Shannon’s entropy indicates more STs with a more even distribution). The results showed that serotypes 6A, 6B, 19F, 16F, and 19A had relatively high ST diversity, while serotypes 5, 38, 7F, 8, and 12F had relatively low ST diversity (Fig. 6C), with ST diversity showing low correlation with sample size (Fig. 6D). These results indicate significant differences in ST diversity among individual serotypes.

**Fig. 6.**
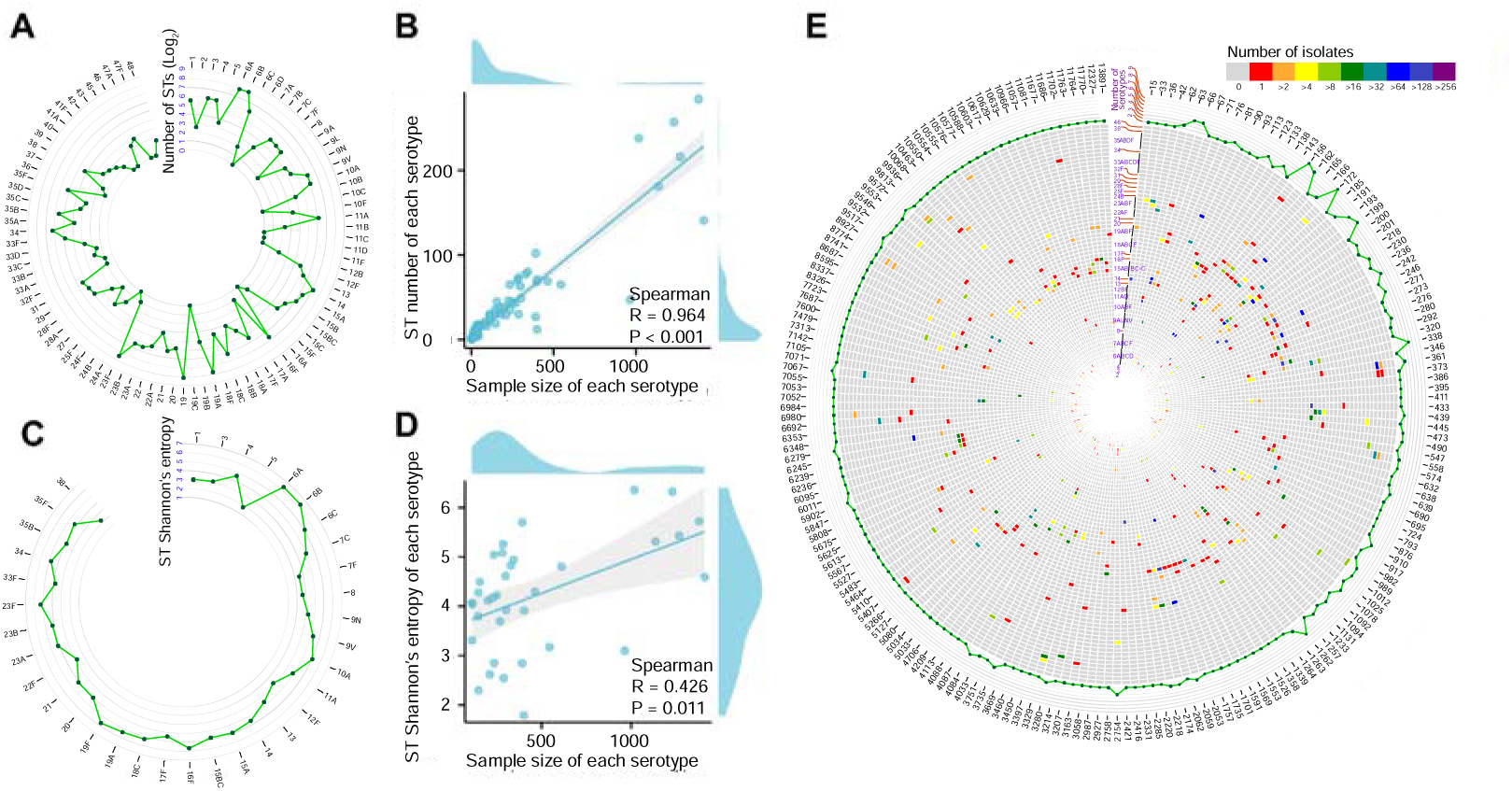
Serotype and ST associations. **A** Number of ST types associated with each serotype. **B** Correlation between sample size and ST type number of each serotype. **C** Shannon’s entropy of STs associated with each serotype. **D** Correlation between sample size and ST Shannon’s entropy of each serotype. **E** Number of serotypes (two or more) associated with each ST. Progressing from the innermost to the outermost part, rings 1 through 70 present heatmaps depicting the count of isolates per serotype for each ST, while ring 71 illustrates the total number of serotypes corresponding to each ST.

Conversely, each of the majority—2,618 out of 2,815 STs was linked to a unique serotype. For instance, all isolates of certain STs, such as the 458 isolates of ST217 and the 110 isolates of ST3081, were exclusively associated with serotype 1. Therefore, an isolate’s ST can often help determine its serotype. We selected the STs of the misannotated isolates (Fig. 3) from and found that, except for ST2377 (not found in Table S9), the serotypes associated with these STs (Table S9) did not match the recorded serotypes, but they aligned with the predictions made by Pneumo-Typer (Table 3). It is important to acknowledge that many STs were represented by only a limited number of isolates, indicating the need for more comprehensive sampling to confirm these findings. Besides, a small proportion of STs (197 out of 2,815) were associated with multiple serotypes. Notably, ST156, ST162, ST199, ST172, and ST193 each were related to 10, 9, 8, 7, and 7 serotypes, respectively. Additionally, there were 167 STs associated with two serotypes, 33 with three, 15 with four, and 3 each with five and six serotypes (Fig. 6E). Given that STs represent highly conserved housekeeping genes with generally low mutation rates, the phenomenon of a single ST corresponding to multiple serotypes is likely due to capsular switch events [53].

**Table 3.**
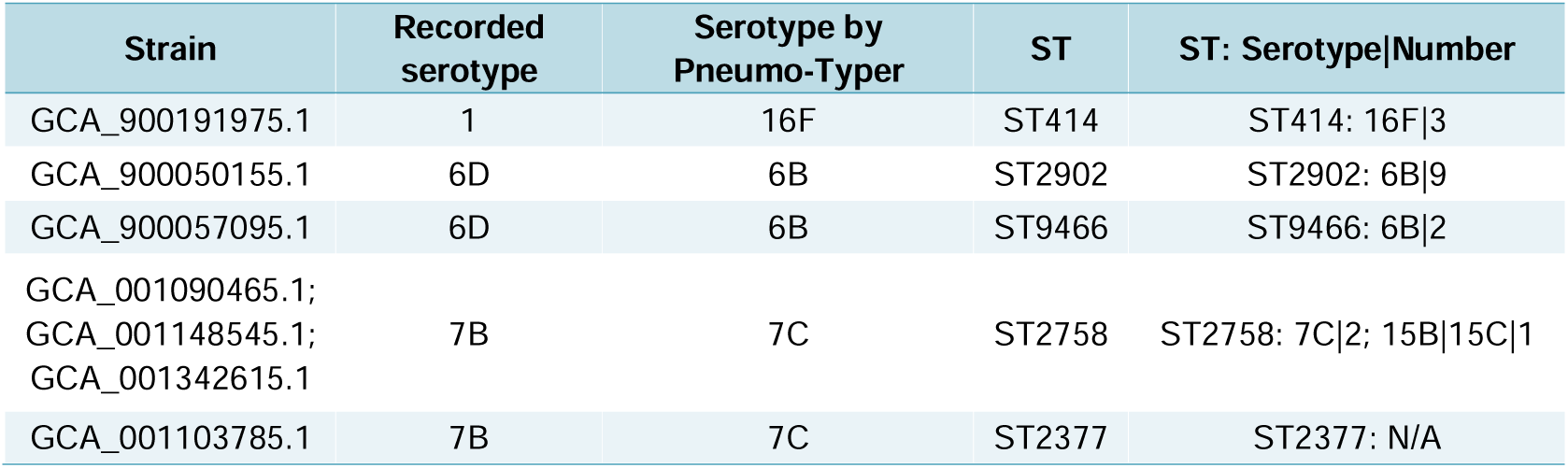
The role of STs in serotype determination.

### Excessive CapT diversity across serotypes

We analyzed CapT diversity across serotypes (Table S10) and found that the number of CapT types varied among serotypes (Fig. 7A). However, as expected, the number of CapT types was highly correlated with sample size (Fig. 7B), indicating that raw counts alone do not accurately reflect true CapT diversity. To address this, we excluded serotypes with fewer than 100 samples and applied Shannon’s entropy to assess CapT diversity. The results showed that serotypes 6A, 7F, 23F, 6C, and 23B had relatively high CapT diversity (Fig. 7C), with no correlation to sample size (Fig. 7D). These findings highlight significant differences in CapT diversity among serotypes, suggesting that different serotypes undergo capsule genetic modifications at varying frequencies. Furthermore, CapT diversity and ST diversity reflect changes in distinct genomic regions, showing only a weak correlation (Fig. 7E). This suggests that CapT and ST evolution occur independently rather than in a synchronized manner.

**Fig. 7.**
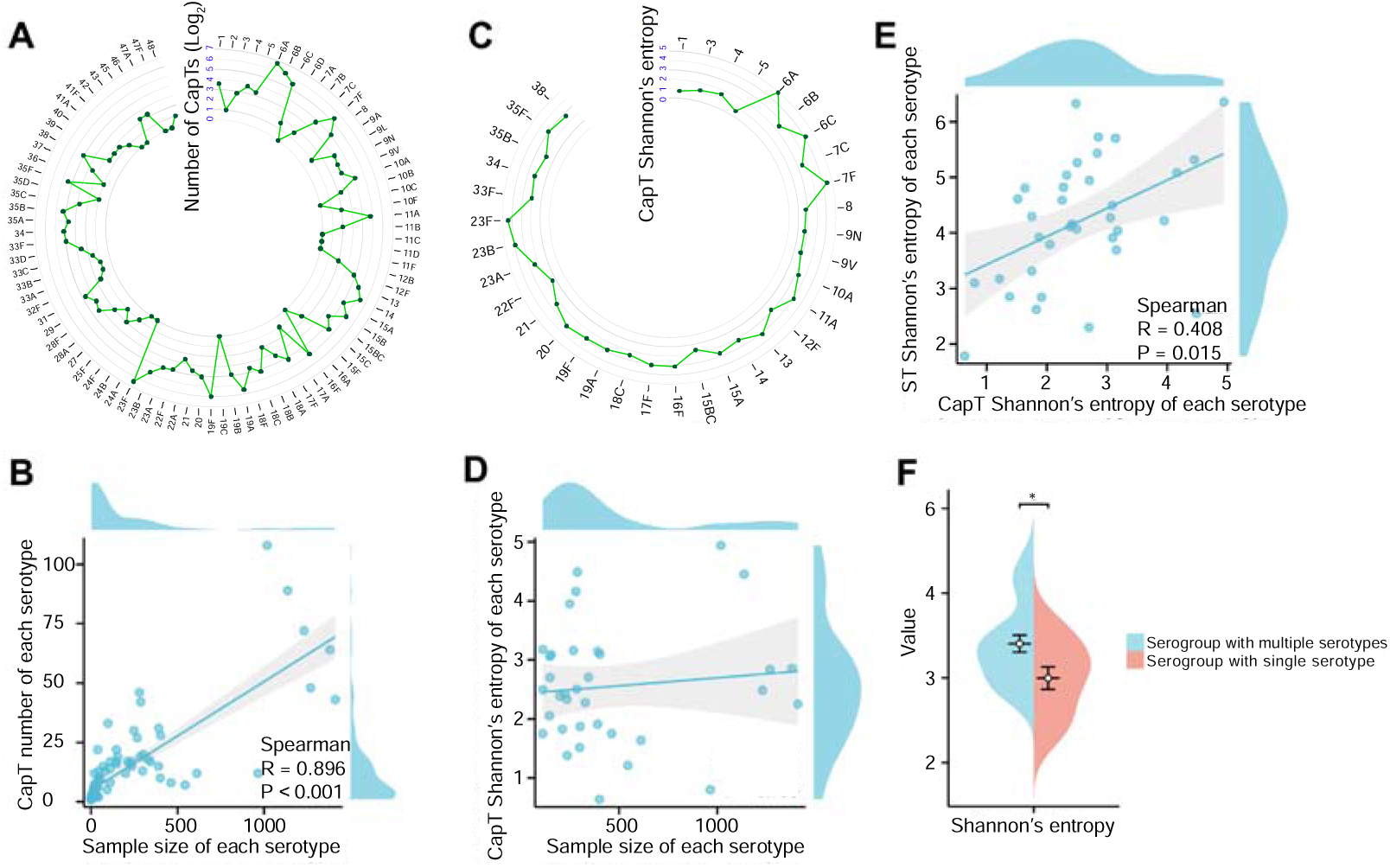
CapT diversity across serotypes. **A** Number of CapT types associated with each serotype. **B** Correlation between sample size and CapT type number of each serotype. **C** Shannon’s entropy of CapTs associated with each serotype. **D** Correlation between sample size and CapT Shannon’s entropy of each serotype. **E** Correlation between CapT Shannon’s entropy and ST Shannon’s entropy of each serotype. **F** Difference in CapT Shannon’s entropy between serogroups with multiple serotypes and serogroups with single serotype.

Additionally, serotypes belonging to serogroups with multiple serotypes (e.g., 6A, 7F, and 23F) exhibited higher CapT diversity compared to those in serogroups that contain a single serotype (e.g., 5, 1, and 3) (Fig. 7F). This may be attributed to certain serotypes undergoing more frequent genetic changes in their capsules, with dominant CapT variants gradually emerging as new subtypes within the serotype or serogroup. Notably, serotypes 20 (ranked 6th) and 13 (ranked 7th) also displayed relatively high CapT diversity. As previously discussed, serotype 20 has been further classified into subtypes 20A, 20B, and 20C (Table 2). Regarding serotype 13, we found that in 227 out of 267 strains, the *whaG* gene, which encodes a glycosyltransferase, was functional despite being recorded as a pseudogene in standard databases. However, to date, serotype 13 has not been formally subdivided into distinct subtypes.

### CapT-ST relationships and implications for vaccine design

As mentioned earlier, STs and CapTs evolve independently. Nevertheless, we aimed to identify potential patterns in the relationships between CapTs and STs (Table S11). To visualize these relationships, we used a Sankey diagram to illustrate the associations between CapTs and STs for PCV20-covered serotypes (see Supplementary files). Taking serotype 11A as an example, we found that STs within CC200 and CC580 corresponded to SPC11F/XX/XX, STs within CC33, CC503, CC1079, and NA (unknown) corresponded to SPC11D/XX/XX, while STs in other CCs corresponded to SPC11A/XX/XX (Fig. 8A). This suggests that STs with closely related gene sequences within the same clonal complex (CC) tend to be associated with similar CapTs, indicating the presence of vertical inheritance constraints within specific STs or CCs.

**Fig. 8.**
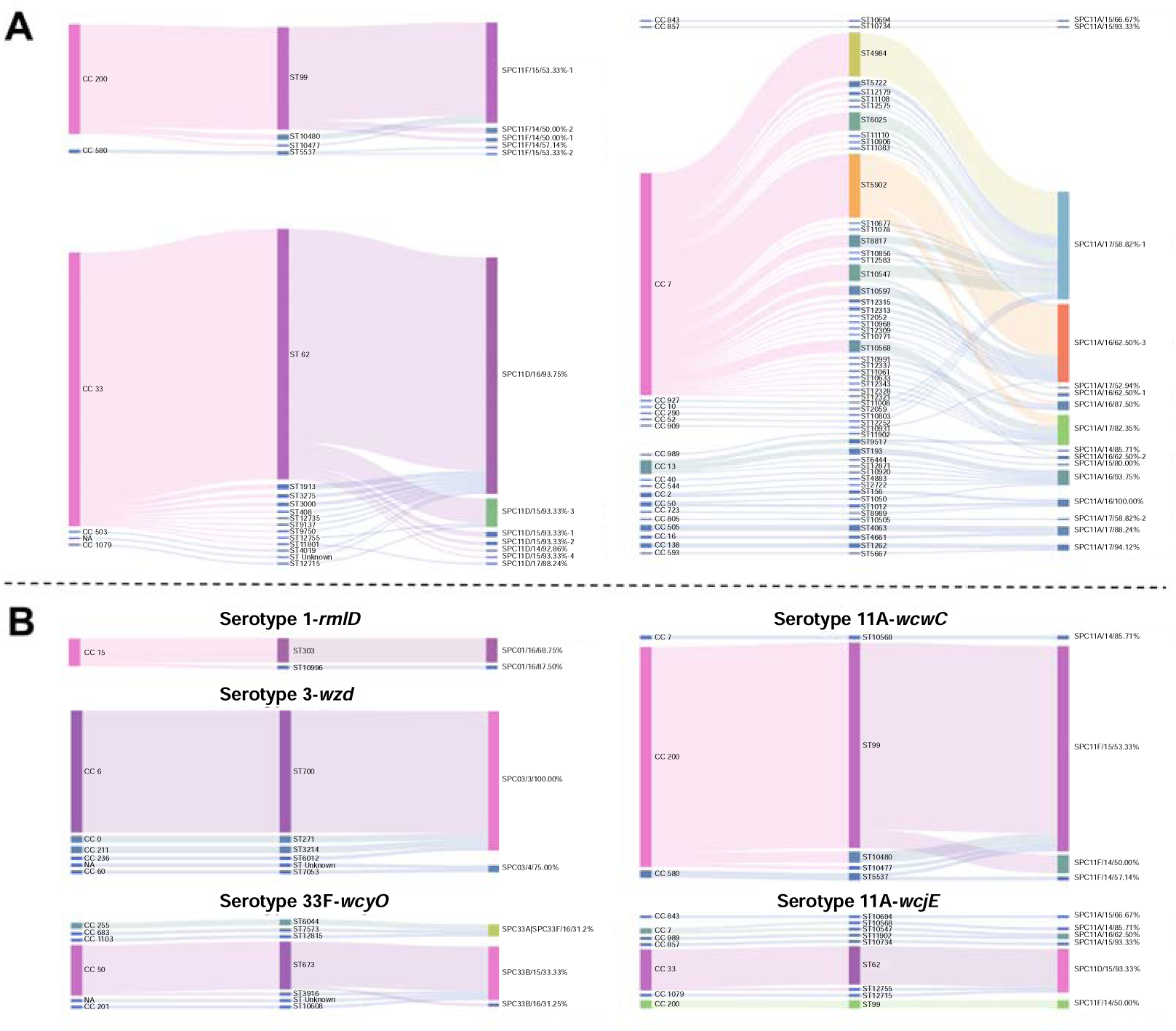
Sankey diagram illustrates the relationship between CapTs and STs for **A** serotype 11A and **B** several serotypes with CapT variations.

Due to these vertical inheritance constraints, when certain STs become prevalent, it is not only essential to monitor their corresponding serotypes but also to consider their associated CapTs, as this has implications for vaccine design. To explore this further, we selected several CapT variations from Table 2 and mapped the relationships between their CapTs and STs (Fig. 8B). Notably, ST303 (serotype 1-*rmlD*), ST700 (serotype 3-*wzd*), ST673 (serotype 33F-*wcyO*), ST99 (serotype 11A-*wcwC*), and ST62 (serotype 11A-*wcjE*) stood out as significant. When these STs are prevalent, it becomes crucial to consider CapT variations in vaccine development. For instance, serotype 3-ST700 exhibits a capsule variant phenotype and has demonstrated enhanced vaccine escape potential following PCV13 introduction [54]. In our dataset, all ST700 isolates lacked the *wzd* gene. Additionally, serotype 11A-ST99 was prevalent in South Korea between 2004 and 2013, accounting for 27.8% of all serotype 11A isolates [55]. In our study, we found that all ST99 isolates carried mutations in the *wcwC* gene. Furthermore, during 2014/15–2018/19 in England, the major circulating STs of serotype 33F were ST717, ST100, and ST673 [56]. In our study, all ST673 isolates contained an insertion of the *wcyO* gene. Therefore, to effectively target infections caused by ST700, ST99, or ST673, it is crucial to incorporate the corresponding CapT variants into vaccine strain selection during vaccine development.

## Discussion

For a long time, epidemiological surveillance of *S. pneumoniae* has primarily focused on clinical outcomes, age distribution, geographical variations, colonization or infection sites, and vaccination status. Further studies have commonly involved analyzing antibiotic resistance patterns and the distribution of serotypes and STs (or cgST, GPSC) among isolates. In recent years, the increasing occurrence of vaccine escape driven by GapT variations has underscored the need for reliable CapT detection. In response, Pneumo-Typer was developed to enable the simultaneous identification of serotype, ST (with optional cgST), and CapT in *S. pneumoniae*.

CapT serves as the genetic basis of serotype determination, while serotypes represent these genetic characteristics at the antigenic level. However, due to significant CapT diversity within individual serotypes, it is no longer feasible to directly infer CapT from serotype alone. CapT is particularly critical for vaccine design, especially in the development of glycoconjugate vaccines, as it provides key insights into capsular antigenic structures. Before the advent of Pneumo-Typer, CapT analysis required a labor-intensive multi-step bioinformatics workflow involving multiple software tools. Pneumo-Typer streamlines this process by integrating visualization features that illustrate capsule gene distribution and operon organization, offering an intuitive understanding of CapT variations.

In the case illustrated in Fig. 4, three isolates were identified as serotype 14 by both SeroBA and SeroCall. While Pneumo-Typer classified two of these isolates as serotype 14 (with another classified as serotype 13/20), its CapT visualization feature revealed substantial differences in the *cps* loci of these isolates compared to the standard serotype 14 strain. Additionally, Pneumo-Typer’s operon organization map identified potential insertions of unknown sequences (depicted as gaps, as seen in PMP1514), prompting further investigation into the origins of these sequences. The inclusion of CapT data thus addresses the limitations of conventional molecular methods for serotype prediction. By leveraging Pneumo-Typer’s CapT analysis capabilities, we identified multiple CapT variations within vaccine-covered serotypes, providing valuable insights for future vaccine design.

Beyond its strengths in CapT detection, Pneumo-Typer also demonstrates robust performance in traditional serotyping and sequence typing. For serotyping, it employs a novel two-step sequence alignment strategy. Unlike PneumoCaT, which relies on full-genome alignment and struggles to determine serotypes for genomes with less than 90% capsular operon sequence coverage, Pneumo-Typer requires only a match to a single capsule gene from its database to assign a serotype. This makes Pneumo-Typer more inclusive, accommodating genomes with incomplete *cps* loci or varying degrees of CapT variation. Despite its inclusivity, Pneumo-Typer maintains high accuracy (98.44%), outperforming other serotyping tools such as PneumoKITy (69.01%) and PfaSTer (90.29%). It is important to note that since SeroBA and SeroCall only support raw sequencing reads, whereas Pneumo-Typer, PneumoKITy, and PfaSTer operate on assembled WGS data, a direct comparison with SeroBA and SeroCall was not conducted in this study. However, we plan to incorporate a raw-read analysis mode in future versions of Pneumo-Typer.

Furthermore, this study represents the first application of CapT and ST data in *S. pneumoniae* serotype determination. Annotation discrepancies that lead to serotype mispredictions can be corrected using Pneumo-Typer’s CapT visualization features. Additionally, ST data provide valuable support for serotype determination, as each ST is predominantly associated with a specific serotype.

The integration of serotype, ST, and CapT data in Pneumo-Typer has enabled a comprehensive analysis of their relationships, revealing several key insights. First, a single serotype can correspond to multiple STs, and the degree of ST diversity varies across serotypes. This observation aligns with prior findings that individual serotypes often exhibit considerable genetic diversity among isolates [57]. Conversely, a single ST is typically associated with only one serotype, reflecting the tendency for capsular genes to spread horizontally over long evolutionary timescales, while clonal expansion with limited serotype switching occurs in the short term [58]. Second, different serotypes display varying degrees of CapT diversity, with multi-serotype serogroups showing higher CapT diversity compared to single-serotype serogroups. This finding is consistent with previous research indicating that multi-serotype serogroups undergo frequent within-group serotype switching events [59]. Lastly, our analysis highlights the relative evolutionary independence between STs and CapTs. This can be explained by the distinct selective pressures acting on core genomes (represented by STs) and non-core genomes (represented by CapTs): STs evolve conservatively under stabilizing selection, whereas CapT variations occur adaptively in response to external pressures such as vaccination or immune evasion [60–63].

## Conclusions

In summary, Pneumo-Typer provides a rapid, accurate, and integrated approach for analyzing serotypes, STs, and CapTs, significantly contributing to epidemiological surveillance and vaccine design for *S. pneumoniae*. Its user-friendly design makes it an excellent tool for researchers. Moving forward, we plan to develop an online version of Pneumo-Typer to further enhance accessibility and usability, particularly for researchers without bioinformatics expertise.

## Availability and requirements

Project name: Pneumo-Typer

Project home page: https://www.microbialgenomic.cn/Pneumo-Typer.html or

https://github.com/Xiangyang1984/Pneumo-Typer

Operating system(s): Platform independent

Programming language: Perl

Other requirements: None

License: GNU General Public License v3.0 or any later version (GPL-3.0-or-later)

Any restrictions to use by non-academics: None

## Abbreviations

CapT: Capsule gentoype
cgST: core genome sequence type
CTVD: Capsular Typing Variant Database
GPSCs: Pneumococcal sequence clusters
IPD: Invasive pneumococcal disease
PCV: Pneumococcal conjugate vaccine
ST: Sequence type
WGS: Whole genome sequencing

## Declarations

### Ethics approval and consent to participate

This study only utilizes data that has been previously published or publicly stored in NCBI database.

### Consent for publication

Not applicable

### Availability of Data and materials

Pneumo-Typer is freely available at https://github.com/Xiangyang1984/Pneumo-Typer and https://www.microbialgenomic.cn/Pneumo-Typer.html. Pneumo-Typer can also be installed through conda: “conda install -c bioconda pneumo-typer”. A detailed README is included. The entire sets of CapT visualization figures of 17,683 pneumococcal genomes and 93 standard serotypes, together with Sankey diagram for PCV20-covered serotypes, are available in Supplementary files. Additional data related to this paper may be requested from the authors.

### Authors’ contributions

ZM and BL conceived and supervised research. ZM and XL devised the data analysis strategies and directed the data analysis. XL implemented the software program while HZ and YZ performed the data analysis and prepared the figures and tables. ZM, ZY, XW, GZ, XZ and YH contributed to the writing of the manuscript. All of the authors have read and approved the final manuscript.

## Acknowledgements

The authors thank Dr. Yingying Hao from the Shandong Provincial Hospital Affiliated to Shandong First Medical University for her needs and feedback regarding the use of the Pneumo-Typer software.

